# Altered Dopamine-Linked Striatal Hemodynamic Latency in Early Psychosis

**DOI:** 10.64898/2026.09.23.753620

**Authors:** Ranesh Mopuru, Blake L. Elliott, Linda J. Hoffman, Lauren M. Ellman, Ingrid R. Olson, Ian C. Ballard

## Abstract

**Importance:** Psychosis is strongly associated with striatal dopamine dysfunction, but noninvasive measures that capture variation in dopamine remain limited. Hemodynamic latency, which quantifies the timing of low-frequency blood oxygen level-dependent (BOLD) fluctuations, may provide a way to indirectly probe dopamine dysfunction in psychosis.

**Objective:** To determine whether striatal hemodynamic latency differs between individuals with early psychosis (EP) and healthy control (HC) participants and whether latency is associated with positive or negative symptom severity.

**Design:** Cross-sectional analysis of resting-state functional magnetic resonance imaging (fMRI) data from the Human Connectome Project (HCP)-EP dataset.

**Setting:** Multisite study at 4 sites using 3T Siemens Prisma MRI scanners.

**Participants:** 105 individuals with EP and 55 HC participants with usable resting-state fMRI and hemodynamic latency data.

**Exposure:** EP status.

**Main Outcomes and Measures:** Voxel-wise hemodynamic latency was estimated using RapidTide and summarized within 6 striatal regions: nucleus accumbens core and shell, anterior and posterior caudate and putamen. Group differences were evaluated using linear mixed-effects models accounting for region and subject-level random effects and controlling for age, sex, and mean framewise displacement. Positive symptoms were measured using the Positive and Negative Syndrome Scale positive subscale; negative symptoms using the Clinical Assessment Interview for Negative Symptoms. Associations with symptoms were assessed using nested linear regressions within the EP group.

**Results:** Latency was significantly lower in the dorsal than ventral striatum in HC and EP, with a greater difference in EP (*b=*-119.46 ms, SE=52.16, *p=*.022). Follow-up six-subregion models found significant main effects of group *F*(1,156.55)=7.79, *p*=.006, and region *F*(5,2760)=187.99; *p*< .001, but no group-by-region interaction *F*(5,2760)=1.74; *p*=.121. Region-specific contrasts showed lower latency in EP in the anterior caudate (*p*=.014), posterior caudate (*p<*.001), anterior putamen (*p*=.007), and posterior putamen (*p*=.029), but not NAc core or shell. Striatal latency was not significantly associated with positive or negative symptom severity among participants with EP.

**Conclusions and Relevance:** EP was associated with broadly reduced striatal hemodynamic latency, with the strongest differences in the caudate and putamen. Altered striatal latency in psychosis warrants further evaluation as a noninvasive indirect marker of striatal pathophysiology linked to dopamine.

**Key Points:** *Question:* Is striatal hemodynamic latency, a novel indirect correlate of dopamine physiology, altered in individuals with early psychosis?

*Findings:* In this cross-sectional study of 105 individuals with early psychosis and 55 healthy comparison participants, early psychosis was associated with significantly lower striatal hemodynamic latency. Differences were observed in the dorsal striatum (caudate and putamen), whereas ventral striatal (nucleus accumbens) latency did not significantly differ between groups.

*Meaning:* These findings support further evaluation of striatal hemodynamic latency as a noninvasive, indirect marker of dopamine-linked physiology in early psychosis

## Introduction

Dopamine is central to biological models of psychosis, with molecular imaging demonstrating elevated presynaptic dopamine synthesis and release in schizophrenia [1]. Although early models emphasized dopamine dysfunction in the ventral striatum, more recent work has increasingly implicated dorsal striatal regions in relation to positive symptoms [2]. Noninvasive measures of regional striatal physiology may therefore provide insight into dopamine-related abnormalities in psychosis.

Functional MRI (fMRI) BOLD signals reflect both neural activity and vascular physiology. This is particularly relevant in neuropsychiatric and pharmacological contexts, where physiological arousal, vascular dynamics, and neurotransmitter systems may influence fMRI signals [3,4]. Systemic low-frequency BOLD fluctuations, often treated as nuisance variance, may therefore contain clinically meaningful information [3,4]. Dopamine may influence these signals because dopamine-receptor occupancy and pharmacological manipulation are linked with hemodynamic responses in dopamine-rich striatal regions [4,5].

Hemodynamic latency captures the relative timing of low-frequency BOLD fluctuations across brain regions, rather than the amplitude of task-evoked activation or resting-state connectivity [6]. Recent evidence shows that striatal latency differentiates nucleus accumbens (NAc) from caudate and putamen, relates to positron emission topography (PET) measures of dopamine physiology, is altered by dopaminergic pharmacological manipulation, and differs in cocaine use disorder [4]. Dopamine released from striatal terminals may act directly on receptors expressed by microvessels, thereby modulating vascular tone [7]. Regional dopamine dysfunction in psychosis could therefore produce altered vascular modulation and shifts in hemodynamic latency. Latency in the striatum may thus offer a novel, non-invasive method of characterizing dopamine-linked changes in vascular physiology (Figure 1). Hemodynamic latency is also influenced by vascular and neuromodulatory factors unrelated to dopamine and therefore should not be interpreted as a dopamine-specific measure. Nevertheless, this metric offers an opportunity to characterize physiological alterations in early psychosis that may be linked to dopamine dysfunction.

**Figure 1.**
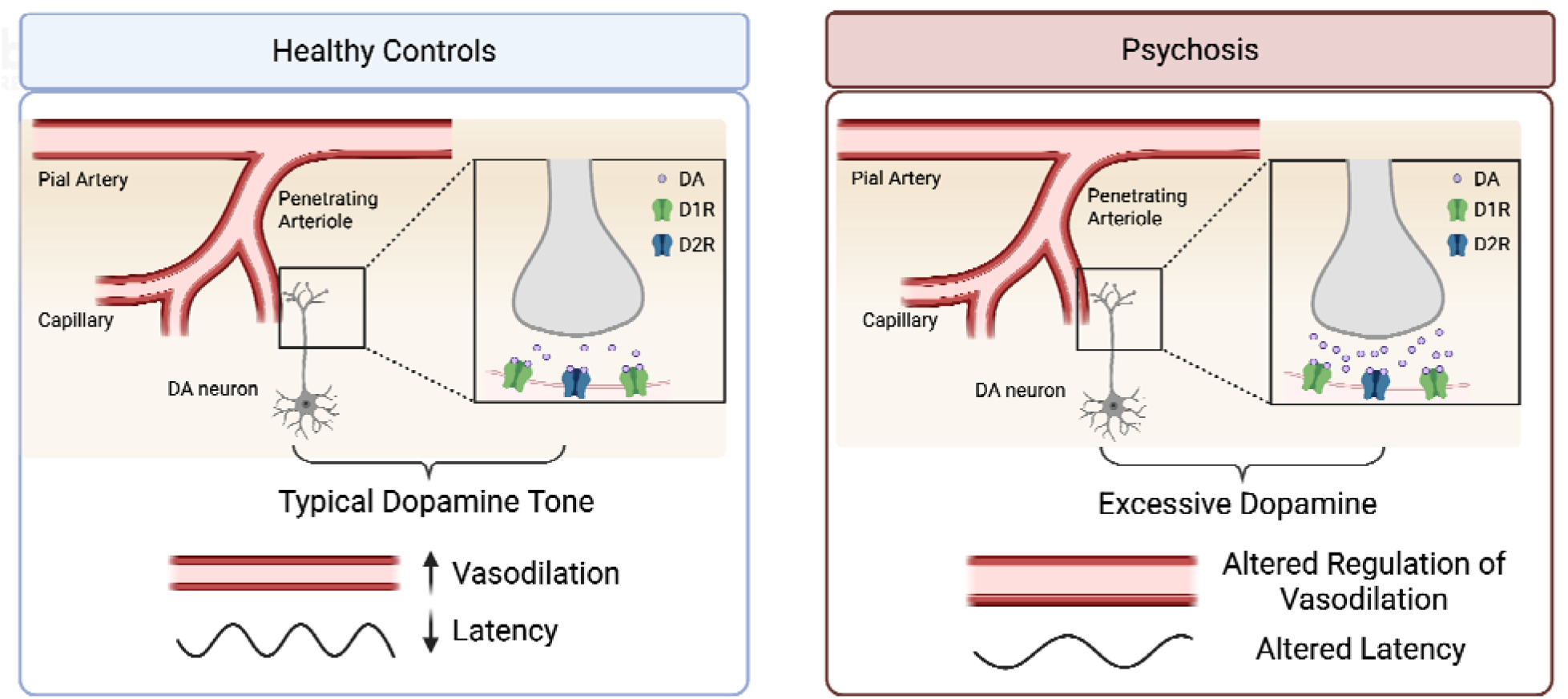
Diagram of the theoretical mechanism through which dopamine may be related to hemodynamic latency (adapted from Ballard et al., 2025 [4]), and how altered dopamine function in psychosis could result in altered striatal latency.

**Figure 2.**
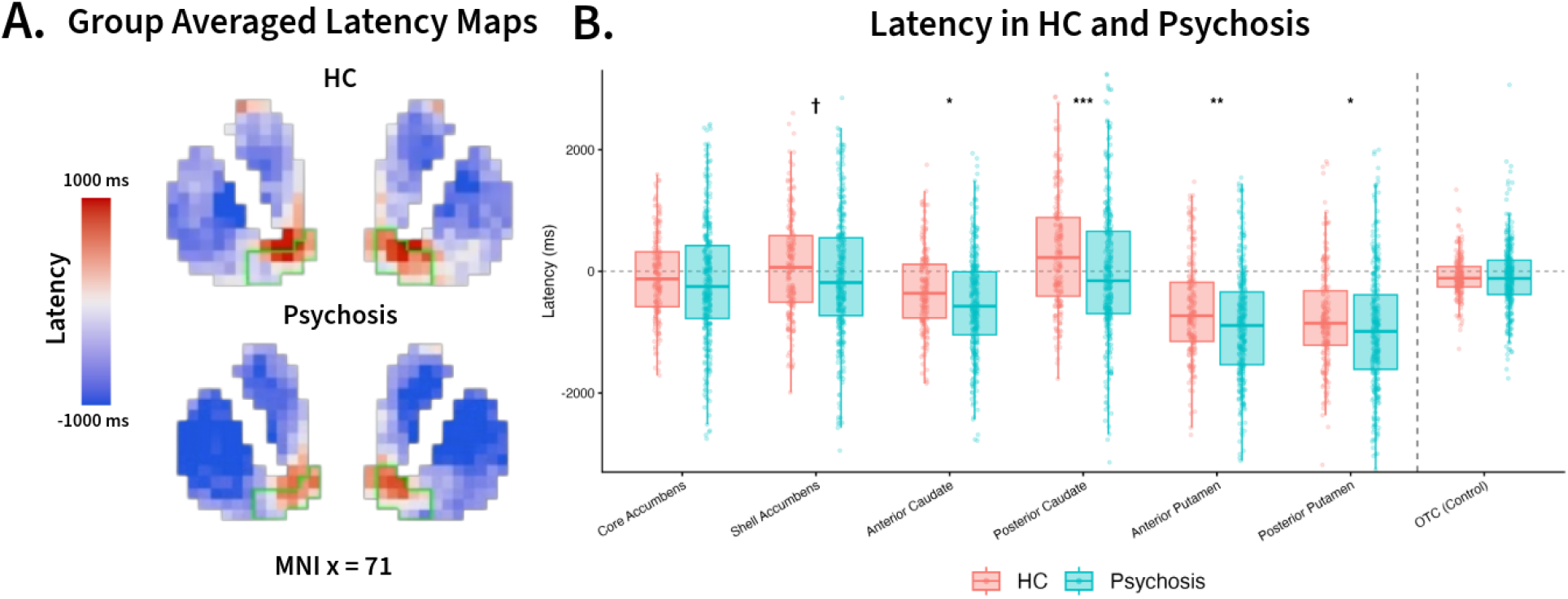
**A**. Group averaged latency maps across the striatum in MNI space for healthy controls (HC) and early psychosis participants. The nucleus accumbens is outlined in green. For ease of visualization, the color range was set to -1000 to 1000 ms. **B**. Boxplots show subject-level mean latency across retained runs for HC and early psychosis participants across six striatal regions and the occipitaltemporal cortex (OTC) control region. Individual points represent participants; boxes indicate the interquartile range with the median shown by the horizontal line. The dashed horizontal line indicates 0 ms latency. P-values are displayed for regions showing significant group differences. † p < .10; * *p* < .05; ** *p* < .01; *** *p* < .001

We tested whether striatal hemodynamic latency differed between individuals with early psychosis (EP) and healthy comparison (HC) participants and whether latency was associated with symptom severity.

## Methods

### Participants and MRI acquisition

Data from 125 EP and 58 HC individuals from the Human Connectome Project for Early Psychosis were analyzed (see Supplement for inclusion criteria). Participants provided written informed consent. Resting-state fMRI data (3T Prisma, two 6-minute runs; AP/PA; TR = 800 ms; 2-mm isotropic resolution) was preprocessed using fMRIPrep. Runs with excessive motion (mean framewise displacement [FD] >0.3 mm or ≥15 frames with FD >1 mm) were excluded.

### Clinical Measures

Clinical assessments were available only for EP participants. Positive symptoms were assessed using the Positive and Negative Syndrome Scale (PANSS) positive symptom subscale [8], and negative symptoms using the Clinical Assessment Interview for Negative Symptoms (CAINS) [9].

### Hemodynamic latency estimation

RapidTide was used to estimate the voxel-wise temporal delay of low-frequency BOLD fluctuations—termed hemodynamic latency—in each useable run, while regressing motion-related nuisance covariates [10]. A gray matter mask was used to generate the global low frequency reference signal.

### Regions of interest and quality control

Striatal regions of interest (ROIs) were defined using the Melbourne subcortex atlas S2 parcellation eroded by a voxel for anatomical specificity, including shell and core nucleus accumbens (NAc), and anterior/posterior caudate and putamen [11]. An occipitaltemporal cortex (OTC) ROI served as a control region [12]. Median latency was extracted from each ROI. Runs with latency estimates exceeding ±3 SD from the sample mean were excluded across three quality-control iterations. Participants were retained if ≥2 resting-state runs remained after quality-control, yielding 105 EP and 55 HC.

### Statistical Analyses

Group differences in striatal latency were tested using linear mixed-effects models with group, region, and their interaction as fixed effects, subject as a random intercept, and standardized age, sex, and mean FD as covariates. A ventral-dorsal model was tested first, followed by a six-subregion model. Type III ANOVA tested fixed effects, and estimated marginal means were used for pairwise group contrasts. OTC latency was analyzed separately using a covariate-adjusted linear regression model. Effect sizes were computed as the group contrast divided by residual SD (Cohen’s *d*; [13]).

Within EP participants, nested linear models tested whether striatal latency collectively predicted symptom severity beyond age, sex, motion, and antipsychotic exposure by comparing covariate-only models with models including all striatal ROIs.

## Results

Latencies were lower in dorsal relative to ventral striatum in both HC (*b=*-342.30 ms, SE=41.42, *p*<.001) and EP (*b=*-461.76 ms, SE=31.71, *p*<.001). This difference was 119.46 ms larger in EP than HC (*b=*-119.46 ms, SE=52.16, *p=*.022). Follow-up six-subregion models showed main effects of group, *F*(1,156.55)=7.79, *p=*.006, and region, *F*(5,2760)=187.99, *p<*.001, but no Group × Region interaction, *F*(5,2760)=1.74, *p=*.121.

Across regions, EP showed lower latency than HC (*b=*-229.42 ms, SE=82.19, *p=*.005, *d* =-0.34). Region-specific contrasts showed lower latency in psychosis in anterior caudate (*b=*-242.10 ms, SE=98.89, *p=*.014), posterior caudate (*b=*-350.35 ms, SE=98.89, *p<*.001), anterior putamen (*b=*-268.35 ms, SE=98.89, *p=*.007) and posterior putamen (*b=*-216.16 ms, SE=98.89, *p=*.029), but not NAc core (*b=*-114.77 ms, *p=*.246), or NAc shell (*b=*-184.79 ms, *p=*.062; Figure 1).

In OTC, a control ROI with limited dopamine innervation or link to EP [4], latency showed no group difference (*b=*-7.98 ms, SE=56.34, *p=*.888). Post-hoc interaction contrasts tested whether the group-difference in each striatal subregion was greater than the difference in OTC. The difference in latency was significantly greater in anterior caudate (*b=*-215.83 ms, SE=86.98, *p*=.013), posterior caudate (*b=*-324.08 ms, SE=86.98, *p*<.001), anterior putamen (*b=*-242.08 ms, SE=86.98, *p*=.005), and posterior putamen (*b=*-189.89 ms, SE=86.98, *p*=.029) than OTC. Striatal latency was not significantly associated with positive or negative symptoms (see Supplemental Text).

## Discussion

EP showed lower striatal hemodynamic latency and greater dorsal-ventral differentiation than HC, with the largest region-specific differences in the anterior and posterior caudate and putamen.

Our findings build on emerging work suggesting that low-frequency BOLD fluctuations contain physiologically meaningful information beyond conventional measures of BOLD amplitude or functional connectivity [3,6]. In the striatum, hemodynamic latency has been linked to PET indices of dopamine function, dopaminergic pharmacological manipulation, and perseverative behavior in a cognitive control task [4]. Consistent with Ballard et al., we observed lower latency in dorsal relative to ventral striatum in HC; extending this finding, the dorsal-ventral difference was larger in EP than HC [4]. The dorsal-to-ventral difference in striatal latency may reflect regional differences in dopamine-related physiology, including slower extracellular dopamine dynamics in the NAc [14]. The greater dorsal-striatal latency reduction in EP is consistent with PET evidence that presynaptic dopamine abnormalities in EP are concentrated in the dorsal striatum [1].

Contrary to our hypotheses, striatal latency was not associated with positive or negative symptom severity in EP. Latency may therefore reflect a diagnosis-related physiological feature rather than current symptom burden, although associations may be smaller, medication-dependent, or more detectable during tasks that dynamically engage dopamine circuitry than at rest.

Hemodynamic latency is an indirect fMRI-derived measure and cannot be interpreted as a direct index of dopamine signaling [3,4,6]. Our findings are therefore consistent with, rather than definitive evidence for, dorsal-striatal dopamine dysfunction in psychosis. Future work should combine latency mapping with PET measures of dopamine physiology and pharmacological challenge in EP. Longitudinal studies will also be important for determining whether striatal latency abnormalities are stable, illness-stage dependent, or sensitive to treatment. Overall, these findings identify altered striatal BOLD timing as a novel physiological feature of early psychosis and support further evaluation of hemodynamic latency as a noninvasive marker of striatal pathophysiology.

## Figures and Tables

**Table 1.** Demographic Information.

| Characteristic | Healthy Controls | Early Psychosis |
| --- | --- | --- |
| **Age, Years** | 24.62 (3.96) | 23.00 (3.79) |
| **Sex, Male** | 35 (63.6%) | 63 (60.0%) |
| **Race** |  |  |
| African American/Black | 4 (7.3%) | 38 (36.2%) |
| American Indian/Alaska Native | 0 (0.0%) | 1 (1.0%) |
| Asian | 8 (14.5%) | 8 (7.6%) |
| White | 40 (72.7%) | 56 (53.3%) |
| Unknown or more than one race | 2 (3.6%) | 2 (1.9%) |
| **Hispanic Ethnicity** | 9 (16.4%) | 5 (4.8%) |
| **Medication Use, Months** | n/a | 13.14 (15.28) |
| **Medication Use, Chlorpromazine Equivalent Dose, mg** | n/a | 150.48 (227.11) |
| **PANSS Positive Score <sup>a</sup>** | n/a | 11.06 (3.92) |
| **CAINS Score <sup>b</sup>** | n/a | 20.30 (9.91) |

Values are presented as *n* (%) or mean (SD). PANSS, Positive and Negative Syndrome Scale; CAINS, Clinical Assessment Interview for Negative Symptoms. <sup>a</sup> Participants with missing values for the variable of interest = 1. <sup>b</sup> Participants with missing values for the variable of interest = 4.

