## Supplement for "Altered Dopamine-Linked Striatal Hemodynamic Latency in Early Psychosis"

**Supplementary Materials for Altered Dopamine-Linked Striatal Hemodynamic Latency in Early Psychosis**

**Supplementary Methods**

**Inclusion Criteria**

Participants were eligible for enrollment if they were medically stable, 16-35 years of age, and provided consent for their deidentified data to be made available through the Human Connectome Project database. Inclusion criteria varied by diagnostic cohort. For the non-affective psychosis cohort, participants were required to meet DSM-5 criteria for schizophrenia, schizophreniform disorder, schizoaffective disorder, psychotic disorder not otherwise specified, delusional disorder, or brief psychotic disorder, with onset of psychotic symptoms within the preceding 5 years. The affective psychosis cohort included participants meeting DSM-5 criteria for major depressive disorder with psychotic features or bipolar disorder with psychotic features, likewise with illness onset within the preceding 5 years (1).

Individuals were excluded for substance-induced psychosis or psychosis secondary to a medical condition; estimated IQ <70 based on available medical history; HIV infection; or neurological or medical conditions that could substantially influence brain function or cognition. The latter included epilepsy or other seizure disorders, stroke, traumatic brain injury or significant head trauma, prolonged loss of consciousness, and other neurological disorders. Additional exclusions included MRI contraindications such as implanted medical devices, severe substance use disorder within the preceding 90 days (excluding nicotine and caffeine), electroconvulsive therapy within the previous year, active suicidal ideation or a suicide attempt within 30 days of screening, or behavior suggesting a substantial risk of harm to self or others.

**Supplementary Results**

For PANSS positive symptoms, the omnibus latency model was not significant. Adding the six striatal latency predictors to the covariate-only model did not improve model fit, *F*(6, 93) = 0.37, *p* = .895, *N* = 104. The residual sum of squares decreased only slightly from 100.03 in the reduced model to 97.68 in the full model, indicating that the six striatal latency predictors collectively explained very little additional variance in PANSS positive symptom severity after accounting for age, sex, mean FD, and antipsychotic exposure.

For CAINS total symptoms, the omnibus latency model was also not significant. Adding the same six latency predictors to the covariate-only model did not significantly improve model fit, *F*(6, 90) = 1.49, *p* = .190, *N* = 101. The residual sum of squares decreased from 93.70 in the reduced model to 85.23 in the full model. Thus, there was no evidence that the combined striatal latency profile significantly predicted negative symptom severity.

Within the full omnibus models, no individual ROI latency coefficient was significant for either outcome. Individual striatal ROI predictors in the PANSS positive symptom model: core accumbens, β = −.24, SE = .14, *p* = .104, 95% CI [−.52, .05]; shell accumbens, β = .00, SE = .12, *p* = .997, 95% CI [−.24, .24]; anterior putamen, β = −.02, SE = .22, *p* = .936, 95% CI [−.45, .41]; posterior putamen, β = −.06, SE = .18, *p* = .721, 95% CI [−.43, .30]; anterior caudate β = .14, SE = .10, *p* = .166, 95% CI [-.06, .34]. and posterior caudate, β = −.22, SE = .14, *p* = .114, 95% CI [−.48, .05].
